# Development of the Mitochondrial Base Editor Analysis Package (MitoBEAP)

**DOI:** 10.64898/2026.06.02.729539

**Authors:** Christian D. Mutti, Pavel Nash, Pedro Silva-Pinheiro, Michal Minczuk, Lindsey Van Haute

**Affiliations:** Medical Research Council Mitochondrial Biology Unit, University of Cambridge, Hills Road, Cambridge, CB2 0XY, UK

## Abstract

For many years, the genetic manipulation of mitochondrial DNA was largely hampered by inefficient delivery of nucleic acids to mitochondria. However, the development of mitoCBEs, such as mitochondrial cytosine base editors (DdCBEs), which catalyse C•G-to-T•A conversions, and more recently, mitoABEs, such as transcription-activator-like effector (TALE)-linked deaminases (TALEDs) enabling A•T-to-G•C conversion, has transformed this field. Generally, mitochondrial base editors exhibit high on-target efficiency and are straightforward to design and use. Nonetheless, unintended off-target effects cannot be overlooked and should be assessed consistently with each experiment, which can be challenging without specialised bioinformatic expertise. Here, we introduce Mitochondrial Base Editor Analysis Package (MitoBEAP), which, to our knowledge, is the first R package specifically designed to analyse next-generation sequencing data from base-edited mtDNA samples. The package facilitates the analysis of potential off-target effects, offers multiple visualisation options, and allows customisation of graphics and thresholds for calculations. As a proof of concept, this study demonstrates how MitoBEAP can be utilised to measure the efficiency of DdCBE treatment targeting human 12S rRNA, as well as to identify potentially harmful off-target conversions across the mtDNA.

## INTRODUCTION

Each cell contains multiple copies of mtDNA, and pathogenic variants of these molecules can cause mitochondrial diseases in humans. The development of genome engineering in mitochondria has been limited until recently due to the inability to efficiently deliver nucleic acids to mitochondria. Recently, there have been significant advances in mitochondrial genome engineering using sequence-specific DNA-binding proteins (such as TALEs or zinc fingers) linked to nucleases (such as FokI), enabling the targeted degradation of mutant mtDNA (1-3). These proteins are engineered with a mitochondrial-targeting sequence (MTS) to facilitate their import across the mitochondrial inner and outer membranes. While these nuclease-based approaches enable the selective elimination of mutant mtDNA, they do not allow the precise introduction of defined nucleotide changes. This limitation has been overcome with the development of mitochondrial cytosine base editors (mitoCBE). DddA-derived cytosine base editors (DdCBEs) enable C•G-to-T•A conversions with high on-target efficiency via transcription activator-like effector (TALE) DNA-binding domains fused to split variants of the bacterial cytidine deaminase DddA. (4). More recently, adenine base editors (mitoABEs) have been developed, such as TALE-linked deaminases (TALEDs), which enable targeted A•T-to-G•C conversions (5). Subsequent optimisation efforts have further improved the performance of these editors, including the development of strand-selective mitochondrial base editors (mitoBEs) (6,7) and the development of engineered DddA variants (e.g., DddA6 and DddA11 (8)) with enhanced activity and broader sequence compatibility, as well as next-generation editors with improved specificity and reduced off-target activity (9-11).

The development of mitochondrial base editors has enabled reverse genetics approaches in mitochondria. For example, libraries of DdCBE editors have been used to introduce premature stop codons across all mitochondrial protein-coding genes, enabling functional knock-out studies of each gene product (12). In addition, multiple groups have demonstrated the use of DdCBE-mediated editing to generate *in vivo* mitochondrial disease models. These include zebrafish models (13,14), generated through targeted mtDNA base editing, rat models harbouring pathogenic mtDNA mutations (15), and mouse models with heritable or tissue-specific mitochondrial genome modifications (16-19). More recently, improved editor variants, including high-fidelity and activity-enhanced systems, have enabled the generation of disease-relevant mouse models with defined mtDNA mutations and associated phenotypes (11,20).

Despite these advancements in mitochondrial base editing with high on-target efficiency, off-target mutations in mitochondrial genomes have been noted, both *in vitro* and *in vivo*, raising concerns regarding the specificity and efficacy of mitoCBEs and mitoABEs for future therapeutic applications. Consequently, the analysis of mtDNA-wide off-targets has become routine when screening and optimising base editors in mitochondria. The methods and parameters used in the analysis can vary across studies, making it difficult to accurately compare findings. To our knowledge, there is no publicly available and standardised tool for the analysis of base editor off-target analysis in mitochondria, limiting both the application of this technology and its further improvement. To close this gap, we developed the Mitochondrial Base Editor Analysis Package (MitoBEAP), an open-source R package available on GitHub that provides a fully standardised workflow for quantifying on-target efficiency, bystander edits, and genome-wide off-target burden after mtCBE or mtABE treatment, regardless of species. MitoBEAP accepts count tables (e.g., samtools mpileup + VarScan) as input, automatically removes mismatches incompatible with the anticipated technology (C•G→T•A for cytosine editors, A•T→G•C for adenine editors), and provides an option to mask haplotype polymorphisms while preserving informative variants. It generates a comprehensive set of visualisations, including scatterplots, bar-chart summaries, and targeting-window heatmaps, along with Excel workbooks that meet NGS repository deposition requirements. To demonstrate the functionality of the R package, we employed DdCBE gene editing to induce C→T nucleotide modifications at positions m.1559 and m.1561 within the 12S rRNA. These positions are proximal to two functionally significant post-transcriptional modifications, m⁴C1491 and m⁵C1493, which are facilitated by the enzymes METTL15 (16, 17) and NSUN4, respectively (21). Subsequent analysis of the base edits by MitoBEAP demonstrates the package’s capabilities and enables in-depth genetic characterisation of the generated cell lines. Next, we used these genetically characterised cell lines to establish the functional relevance of the 1559 and 1561 C-to-T edits in wild-type cells and in the context of METTL15 inactivation. We demonstrate the importance of the 1559 and 1561 positions for the maintenance of mitochondrial ribosome integrity. Additionally, we show that C-to-T edits at 1559 or 1561 in METTL15 knockout cell lines have a cumulative effect.

## MATERIAL AND METHODS

### Overview of the MitoBEAP flow

MitoBEAP, implemented in an R package, provides a computational tool to analyse the efficiency of mitochondrial gene editing and to identify potential harmful unwanted off-target conversions. As input, MitoBEAP utilises count tables, which can be generated using various tools. Here, we use Samtools mpileup and VarScan, ignoring indels. Fig 1 shows a flow chart of the MitoBEAP pipeline. In brief, after uploading the count tables, non-C-to-T/G-to-A (for mtCBE) or non-A-toG/T-to-C (for mtABE) reading errors will be removed and efficiency (on-target) and off-target effects will be calculated. MitoBEAP then gives the user the option to visualise the data in different formats and to create data tables. The details of the functionalities are described below. MitoBEAP can be downloaded at GitHub (https://github.com/NextGenSeek/MitoBEAP), together with the count tables generated for the analysis of gene editing the 12S rRNA, which were used as an example. It should be noted that MitoBEAP possesses additional features not displayed within this manuscript but accessible through the tool’s vignette.

**Figure 1:**
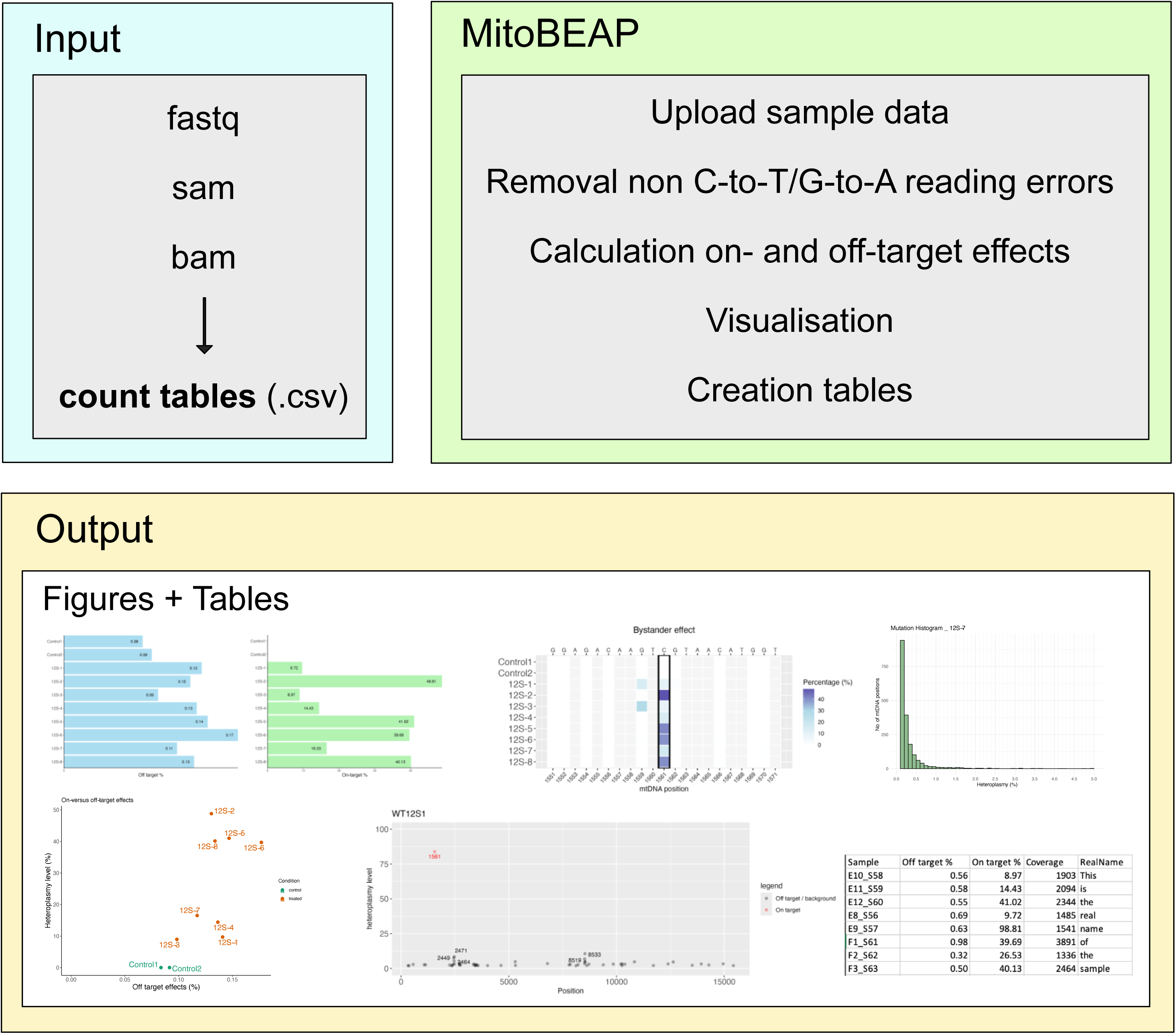
Flowchart showing the principles of MitoBEAP. MitoBEAP requires count tables as input, which can be generated in many ways. The package contains functions to remove reading errors and calculate on- and off-target effects. A wide range of publication-ready visualisation options and functions for generating summary tables provide the user with all the necessary tools to analyse their gene-editing experiments.

### Plasmid construction

The DdCBE architectures employed in this study were as previously reported (4). TALE arrays were designed utilising Repeat Variable Diresidues (RVDs) encompassing NI, NG, NN, and HD amino acids, corresponding to A, T, G, and C, respectively. For construction of the plasmids utilised in cell screens, all DdCBEs ORFs were synthesized as gene blocks (GeneArt, Thermo Fisher). DdCBEs targeting the L-strand of the mtDNA were inserted into a pTracer cytomegalovirus promoter (CMV)/Bsd (pTracer) backbone along with eGFP, while those targeting the H-strand were integrated into a pcDNA3.1(−) mCherry (pcmCherry) backbone accompanied by mCherry expression.

### Cell culture and transfections

Wild-type (WT) and METTL15 KO HEK293T cells were cultured as a monolayer at 37°C in a humidified atmosphere with 5% CO2 using high-glucose DMEM (Gibco) supplemented with 10% foetal bovine serum, 100 U/mL penicillin, 50 μg/mL uridine and 100 μg/mL streptomycin. For screening DdCBE pairs, human HEK293T cells were seeded into 6-well tissue culture plates at approximately 70% confluency and transfected with either 3,200 ng of each monomer (L and H). Transfection was performed with 16 µl of FuGENE-HD (Promega), following the manufacturer’s instructions. After 24 hours, cells were harvested for FACS analysis and sorted for GFP and mCherry double-positive cells using a BD FACSMelody cell sorter. The sorted cells were then allowed to recover for an additional 6 days before DNA extraction, as described below.

### Measurement of cell growth

For cell growth assays, HEK293T cells transfected with DdCBE were cultured in either glucose-containing DMEM (4.5 g/l glucose, 110 mg/l sodium pyruvate, 10% FBS, 100 U/ml penicillin, 100 μg/ml streptomycin) or galactose-containing DMEM (0.9 g/l galactose, 110 mg/l sodium pyruvate, 10% FBS, 100 U/ml penicillin, 100 μg/ml streptomycin). Confluency was assessed using an Incucyte S3 live-cell imaging system (Essen BioScience), capturing measurements at ×4 zoom, with 9 images per well taken every 6 hours.

### Genomic DNA isolation and Sanger sequencing

Cells were harvested by trypsinisation, followed by a single wash in PBS and resuspension in lysis buffer (1 mM EDTA, 1% Tween 20, 50 mM Tris, pH 8) supplemented with 200 µg /ml proteinase K. The lysates were agitated at 56 °C (300 r.p.m.) for 1 hour, then incubated at 95 °C for 10 minutes before use in downstream applications.

For Sanger sequencing, approximately 300 bp regions of 12S rRNA were PCR-amplified using GoTaq G2 DNA polymerase (Promega) and specific primers listed in **Supplementary Table 1**. The PCR protocol involved an initial heating step of 1 minute at 95 °C, followed by 35 cycles of amplification (30 seconds at 95 °C, 30 seconds at 62 °C, 15 seconds at 72 °C), and a final extension step of 5 minutes at 72 °C. PCR purification and subsequent Sanger sequencing were performed by GENEWIZ/AZENTA (UK).

### Library preparation and mtDNA-wide sequencing

For comprehensive mtDNA-wide sequence analysis, two overlapping long amplicons covering the entire mtDNA molecule were generated through long-range PCR with PrimeSTAR GXL DNA polymerase (TAKARA), using the primers as reported in (4).

The PCR protocol began with an initial denaturation step of 1 minute at 94 °C, followed by 16 cycles of amplification (30 seconds at 98 °C, 30 seconds at 60 °C, 9 minutes at 72 °C), and ended with a final extension step of 5 minutes at 72 °c. Subsequent tagmentation and indexing PCR were performed using Nextera XT (Illumina), following the manufacturer’s guidelines. The libraries were then sequenced using high-throughput sequencing on the Illumina MiSeq or NovaSeq platform (PE250/PE150) and demultiplexed with the manufacturer’s software.

### Western blotting

Cell pellets were lysed using lysis buffer (50 mM Tris-HCl pH 7.4, 1 mM EDTA, RNasin Inhibitor, 1% Triton X-100, 1/50 Roche Inhibitor). Lysates were clarified by centrifugation at 10,000 rpm for 5 min at 4°C. The lysates were transferred to clean tubes and kept at -20°C. Protein concentration in clarified lysate was quantified using the PierceTM BCA Protein Assay (ThermoFisher Scientific), following manufacturer’s guidelines. Approximately 30 μg total protein was diluted to an equal volume and combined with NuPAGE LDL Sample Buffer 4× (Invitrogen) and 50mM DTT. The samples were heated to 92°C for 5 min and loaded on SDS-PAGE 4-12% bis-tris gels (ThermoFisher Scientific) at 200 V for 25 min. Proteins in SDS-PAGE gels were transferred to nitrocellulose membranes using iBlot 2 Dry Blotting System (ThermoFisher Scientific). The membranes were blocked for 1 h with 5% milk in PBST at RT, then primary antibody overnight at 4°C followed by secondary for 1 h at RT. The blots were imaged using an Amersham Imager.

## RESULTS

### Targeted cytosine base editing of the 12S ribosomal RNA

To generate data demonstrating the function of MitoBEAP, we chose to use DdCBE to modify the 12S rRNA in the decoding centre of the mitoribosome. We aimed to introduce the m.1561C>T mutation, as this would disrupt RNA base pairing in helix 44, close to two post-transcriptional modified positions (m^4^C1491 and m^5^C1493) (**Supplementary Figure 1A-B**). To identify the most efficient DdCBE pair, we generated 32 combinatory pairings of varying DdCBE 1397 and 1333 splits (**Supplementary Figure 1A-C**), which were first screened by Sanger sequencing (**Supplementary Figure 1D-E**). Eight pairs showed base editing at the designated target position with varying levels of on-target efficiency. The edits produced by these pairs were subsequently analysed using mtDNA-wide next-generation sequencing (NGS) to more precisely assess both on-target and off-target efficiencies.

### MitoBEAP: Efficiency and off-target calculations

According to their website, Illumina sequencing has a 99.9% base call accuracy for a Phred Quality score of 30, yet, due to the large amount of data generated, even such a high accuracy still results in thousands of read errors. Although rare, library preparation can also introduce mistakes as it involves PCR steps. On-target activity of DdCBEs is based on catalysing C•G-to-T•A conversions (4), hence, off-target conversions are also predicted to be limited predominantly to C•G-or-T•A. Therefore, MitoBEAP removes all other mismatch combinations before calculating off-target effects. In this paper, we use the *OnTargetCalcCBE()* function, optimised for C•G-or-T•A conversion, but MitoBEAP also has the function *OnTargetCalcABE()*, specifically optimised for adenine base editing.

Mitochondrial DNA is highly polymorphic, so samples are likely to differ from the reference genome. Since polymorphic differences relative to the reference genome should remain unchanged between treated and control samples, the user might choose to display those polymorphisms and include them in the calculations as background effects. This is the standard option in MitoBEAP (**Supplementary Figure 2A**). However, the user might prefer to exclude such sites from the analysis; therefore, MitoBEAP allows the user to remove measurements for certain positions across all samples if the heteroplasmy levels in the control samples exceed a user-defined minimum heteroplasmy level for that position in the genome. (**Supplementary Figure 2B**). However, this option should be used with caution. Positions that are nearly 100% heteroplasmic in all control samples are often discarded because they are likely to reflect mtDNA haplotypes. Yet, even these sites are not completely shielded from unintended base-editor activity, which can introduce new conversions and distort interpretation if the sites are removed from analysis. Consequently, the exact threshold level may vary between projects. In this study, we used a threshold of 70%.

MitoBEAP generates scatter plots for all C•G-to-T•A conversions, enabling users to quickly screen a preliminary sample for off-target effects. Here, the user also has the option to remove low-level heteroplasmy positions, inevitably present in any sample, for better visualisation of off-target conversions (**Figure 2A**, threshold = 1%). It should be noted that those values will only be removed for visualisation but will still be included in the off-target calculations. In this instance, the 12S rRNA at on-target position 1561 has been modified by approximately 50%. Although there are some off-target modifications, their extent never exceeds 20%.

**Figure 2:**
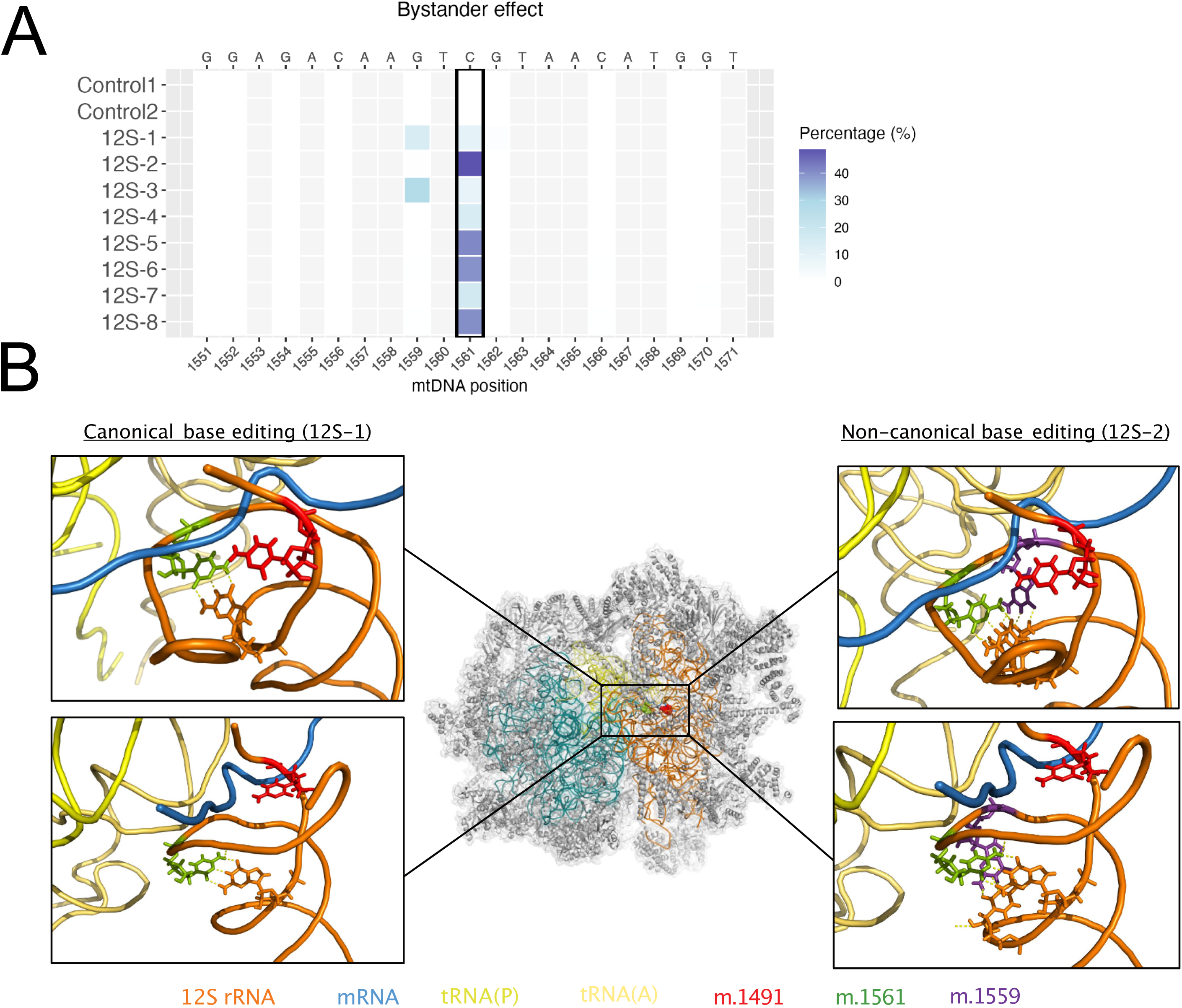
Selection of the optimum construct for DdCBE treatment of human 12S rRNA. (A) Scatter plot showing the efficiency of cells transfected with the 12S-2 DdCBE construct, corrected for high heteroplasmy levels detected in the control samples. Only positions with heteroplasmy levels above 1% are shown. (B) Bar chart providing an overview of the efficiency (on-target effect) of the tested constructs. (C) Bar chart showing the average off-target levels for each tested construct. (D) Overview of the off-target percentage (x-axis) with heteroplasmy levels on the y-axis for all tested constructs (red) and control samples (green).

MitoBEAP’s *CreateBarChart()* function produces bar charts displaying on-target (**Figure 2B**) or off-target (**Figure 2C**) edits, while *AdjScatter()* creates a combined plot for easy comparison between samples (**Figure 2D**).

The MitoBEAP package not only includes several visualisation options but also enables the user to generate overview tables, which allow for quick comparison of samples (**Supplementary Table 1**). Additionally, data repositories such as GEO require the user to upload data tables alongside the raw fastq.gz files. MitoBEAP can generate such tables in Excel format, with a separate worksheet for each sample (**Supplementary File 1**).

### MitoBEAP: Visualisation of bystander editing

In addition to the intended genetic alterations, unintended bystander mutations frequently arise within the base editor’s targeting window (i.e., the sequence between the TALE binding sites of the DdCBE monomers). These mutations are particularly challenging to eliminate entirely; therefore, users are advised to compare different DdCBE or CBA pairs. To facilitate visualisation of bystander editing, mitoBEAP offers an intuitive heatmap feature that enables users to visualise the targeting window (**Figure 3A**).

**Figure 3:**
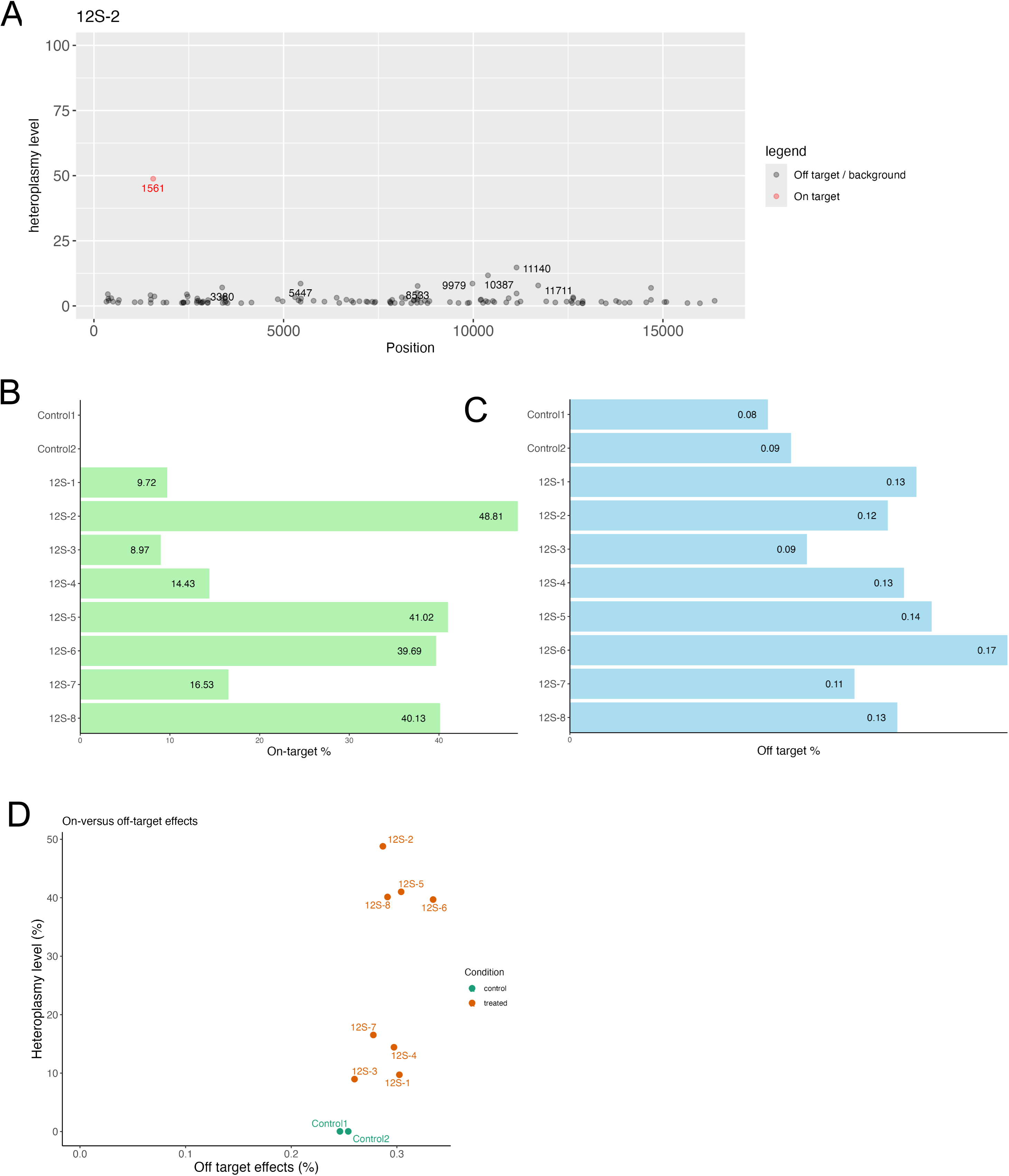
Bystander effect in 12S rRNA following DdCBE treatment. (A) MitoBEAP-generated heatmap illustrating the bystander effect, demonstrating that DdCBE-mediated mitochondrial base editing can target non-canonical 5’-GACT-3’ sites. The rectangle highlights the on-target position (1561) (B) The locations of the canonical and non-canonical base-edited sites within the structure of the mitochondrial ribosome decoding centre. These mutations are adjacent to the region where methyltransferase METTL15 deposits m^4^C (m.1491C), which has been shown to be essential for mitoribosome biogenesis.

In the presented experiment, we observed non-canonical editing outside the 5’-TC sequence context, specifically at 5’-GACT-3’. The non-canonical editing broadens understanding of the positions that can be modified by DdCBEs. This finding also indicates the potential to produce double mutants in a previously uncharacterized context utilising DdCBEs, thereby expanding their scope of application. Within the mitochondrial ribosome, these two mutations are expected to significantly disrupt rRNA base pairing, thereby destabilising the rRNA structure. (**Figure 3B**).

MitoBEAP has many additional functions and options that fall outside the scope of this manuscript, which can be found in the most recent vignette, available on GitHub.

### Assessing the effect of mutating the 12S rRNA on the mitoribosome

All tested constructs exhibited comparable levels of off-target base conversions across the mitochondrial genome, with an average frequency of approximately 0.1%. This value is based on quantifying C:G-to-T:A conversions at all cytosine residues genome-wide, and it is similar to the baseline heteroplasmy observed in untreated control samples. Such low off-target rates indicate the high specificity of these DdCBE pairs for mitochondrial genome editing. (**Figure 2C**). Furthermore, for the selection of optimal DdCBE pairs to target both canonical and non-canonical mtDNA positions, constructs were evaluated based on their ability to provide the highest on-target editing efficiency while maintaining the lowest possible off-target conversion rates (**Figure 2A–E**). This rigorous comparison ensured that only the most precise and efficacious editor pairs were prioritised for downstream applications. (**Figure 2A-E**).

Considering the proximity of the base-edited positions within the 12S rRNA to the METTL15 modification site (22), a further study was conducted to evaluate the impact of editing these positions in METTL15 knockout cells. MitoBEAP was employed to analyse both on-target and off-target activities across the mitochondrial DNA (**Supplementary Figure 3A-C**). High on-target editing levels of 75% and 84% were detected in the METTL15 KO cells, with some presumably insignificant low-level off-target edits. In WT cells transfected with the 12S-2 construct, a heteroplasmy level of 59.4% was observed, while 10.4% C-to-T editing at the m.1559 position was measured in the METTL15 KO (**Supplementary Figure 3D**). These cell lines were subsequently used for further analysis to evaluate the impact of these mutations on the mitochondrial ribosome.

To demonstrate that our tool facilitates the selection of optimal gene editing pairs aligned with the user’s objectives, we further characterised our base-edited cell lines. An evaluation of the effect of base editing of the 12S rRNA on the mitoribosome was conducted via Western blot analysis (**Figure 4 A-B**). The METTL15 KO cell line with the m.1561C>T mutation (75% heteroplasmy) exhibits a significant impact on mitochondrial protein levels. All the probed mitoribosome proteins are almost undetectable and markedly lower than WT levels. The same observation is seen with the OXPHOS subunits, particularly complexes I and IV cannot be detected by western blotting (**Figure 4A-B**). Further analysis of these data indicates that depletion of METTL15 and mutation of the 12S rRNA have a cumulative effect on the mitochondrial ribosome. Individually, the METTL15 KO or 12S-1 (m.1561C>T) mutated HEK293T cell lines show comparable, mild phenotypes in these experiments. However, when both modifications are present, there is a drastic impact on the levels of all mitochondrially encoded proteins that were examined. This finding was confirmed through growth assays in galactose-containing medium, which forces mitochondrial ATP production via OXPHOS. (**Figure 4C-D**). The METTL15 KO cells harbouring the m.1561C>T mutation entirely failed to proliferate in media containing galactose. The data substantiate that mutations in 12S rRNA, combined with the absence of METTL15, which removes the m^4^C modification, have a cumulative and detrimental effect on the mitochondrial ribosome and, consequently, on mitochondrial function. These experiments further indicate that the non-canonical m.1559C>T mutations exert negative effects on mitochondrial health, even at very low heteroplasmy levels, thereby highlighting the importance of maintaining the structural integrity of the 12S rRNA decoding centre in mitochondrial ribosome biogenesis and function.

**Figure 4:**
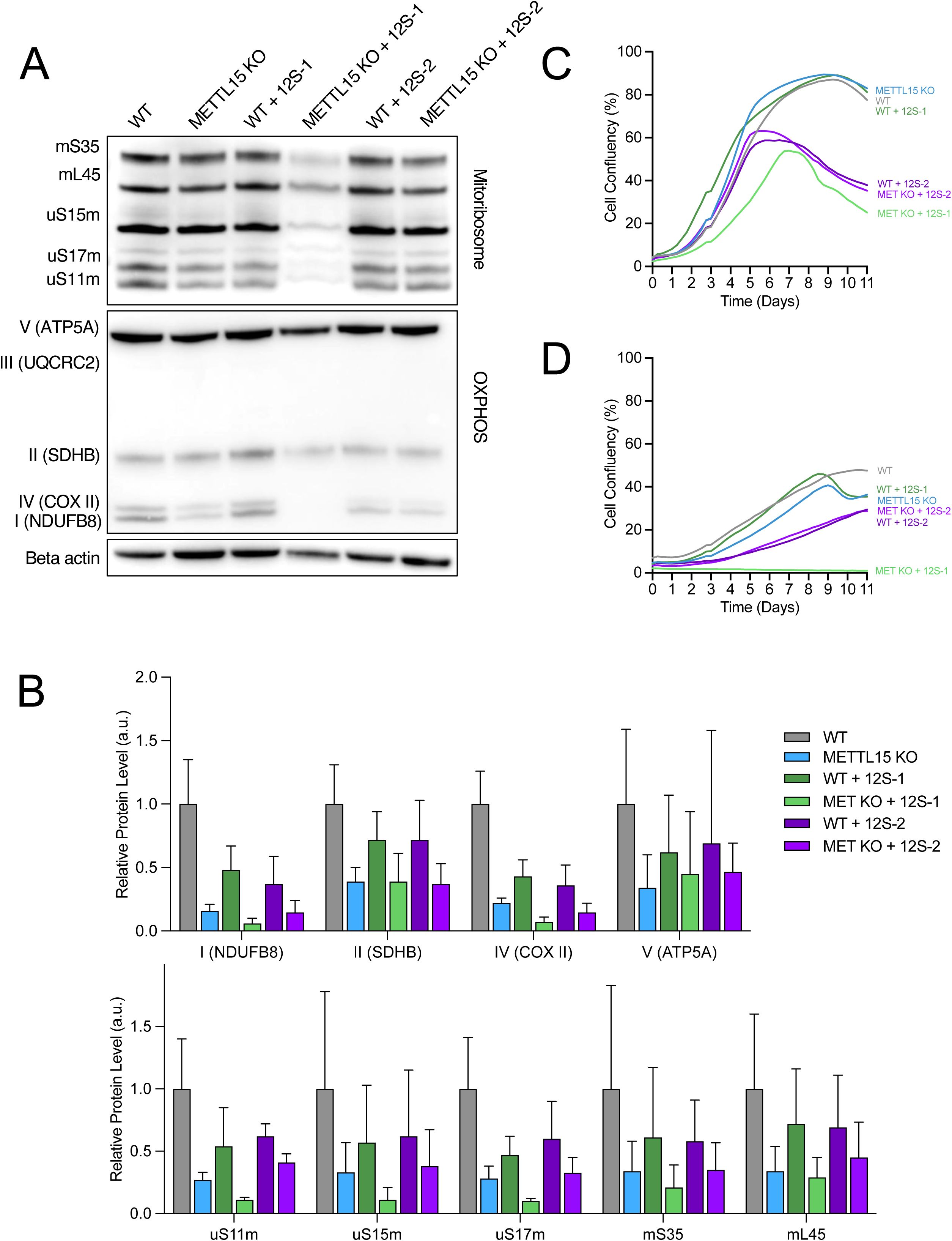
Characterisation of selected DdCBE-generated cell lines. (A) Representative western blotting of proteins in the mitochondrial ribosome and components of the OXPHOS complexes in cells with and without base editing of 12S rRNA. (B) Quantification of blots in (A) using ImageJ, n = 3. (C- D) Growth curves of base-edited cells grown in either glucose (C) or galactose (D).

## DISCUSSION

Following the development of DNA base editing in mitochondria, the possibilities of using a reverse genetics approach to study mitochondrial function has expanded considerably. NGS is widely used to assess the impact of base editors on the mitochondrial genome. MitoBEAP is a user-friendly R package, specifically designed to analyse both the desired base change and off-target edits in mitochondrial base editing. Until now, research groups had to create their own bioinformatic pipelines to analyse mtDNA editing efficiency and off-target levels. MitoBEAP provides researchers with a standardised pipeline, enabling better comparison between experiments.

Given the importance of post-transcriptional modification of rRNA for its structural and functional activity, we employed base editing to mutate the 12S rRNA in the decoding centre of the mitoribosome. The hypothesis was that disrupting RNA base pairing within the decoding centre would have a similar effect to the lack of modification (22), thereby affecting the translating ability of mitoribosomes. Given that the m.1491C position modified by METTL15 is not located within a canonical 5′-TC-3′ context, it was not targetable with the DdCBE architectures available when this study was initiated. Therefore, the adjacent site at m.1561C, which lies within a compatible sequence context, was selected for editing. Conversion of m.1561C>T disrupts an RNA base pairing in helix 44 close to both the m^4^C and m^5^C modified positions (**Figure 3A-B**). By disrupting interactions within this helix, 12S rRNA is unlikely to form its secondary structure correctly, with strong implications for the small subunit of the mitoribosome. During the course of this work, whilst using our newly developed mitoBEAP package, we discovered that some of the screened base editors exhibit non-canonical editing outside the 5’-TC-3’ context, occurring at 5’-GACT-3’ sites. We utilised this to generate double-mutant cell lines in 12S rRNA, alongside single mutants. When combined with the METTL15 knockout cell line from our previous study (22), we were able to demonstrate cumulative effects of the integrity of the mitochondrial ribosome. This work has further reinforced the significance of the decoding region of the 12S rRNA in the biogenesis, stability, and activity of the mitochondrial ribosome. Follow-up studies using cryo-electron microscopy of the mitochondrial ribosome in these cell lines could aid in capturing the assembly pathway state caused by this disruption of 12S rRNA. Given the substantial destabilisation of the 12S rRNA, this may be a challenging state to purify, and there is likely to be considerable heterogeneity of the ribosome in these cells.

Here, we show that the publicly available R package MitoBEAP, which is optimised to analyse and standardise mitochondrial base editing, can be utilised to select the most suitable base editors for a specific target and to evaluate its on-target efficiency as well as potential unintended off-target effects. As proof of concept, we performed DdCBE base editing of the human 12S rRNA and further examined the selected cell lines to confirm its biological relevance.

## SUPPLEMENTARY FILES

Supplementary File 1: MitoBEAP Package Vignette. The most recent version can always be found on GitHub: https://github.com/NextGenSeek/MitoBEAP/tree/main/vignettes

Supplementary Table 1: Primer sequences used in this study

## DATA AVAILABILITY

The data supporting the findings of this study are available within the paper and its Supplementary Information. The NGS files generated in this study are available in the GEO database under accession number GSE309661.

## ACKNOWLEDGEMENTS

We acknowledge the members of the Mitochondrial Genetics Group (MRC-MBU, University of Cambridge) for useful discussions during the course of this research. The FACS experiments were performed by the CIMR Flow facility, and NovaSeq sequencing was performed in the Genomics Facility of the Cancer Research UK (CRUK) Cambridge Institute.

## Supplementary figures

**Supplementary Figure 1: Base editing of the 12S rRNA**

(A) Structure of the mitochondrial ribosome (PDB ID: 6ZSG) showing the decoding centre of the mtSSU. (B) Zoomed boxes show the base edited position m.1561 in green and position m.1491, methylated by METTL15, in red in helix 44 of the 12S rRNA. (C) The architecture of DdCBE monomers targeting the 12S ribosomal RNA decoding site. DNA binding is achieved through TALE domains designed specifically to target the mtDNA L-strand and H-strand of the 12S rRNA gene at positions m.1561C (green) and m.1559G (purple). Cytosine deaminase base editing is performed by the DddAtox, which is split to reduce toxicity and is attached to the 3’ end of each TALE repeat. Once the TALE regions bind to the specific mtDNA, the split DddAtox halves reconstitute to generate an active cytosine deaminase. MTS SOD2 mitochondrial targeting sequence from superoxide dismutase 2, UGI uracil glycosylase inhibitor, L-strand or (L) light mtDNA strand, H-strand or (H) heavy mtDNA strand. (C) Schematic of TALE arrays used in screens designed to target H- and L-strand sequences in the 12S rRNA gene of human mtDNA. Two different TALE domains targeting either L- or H-strand sequences flanking the decoding region of the 12S rRNA gene were combined with N- or C-terminal halves of DddA split at either G1333 or G1397. (D) Example of Sanger sequencing traces used for (E) Quantified heteroplasmy shifts from Sanger sequencing traces of base-edited HEK293T cells, represented as a matrix of DdCBE monomer pair combinations and their base-editing efficiency. Thick black squares in screening matrices indicate the chosen DdCBE monomer pairs (H2-1333-C and L1-N-1333-N).

**Supplementary Figure 2: Example figure showing different thresholds**

(A) Scatter plot created with MitoBEAP displaying all positions for sample 12S-2. The on-target position is marked in red. (B) MitoBEAP-generated figure, adjusted for high heteroplasmy levels found in the control for sample 12S-2. The on-target position is indicated in red.

## REFERENCES

1. Gammage, P.A., Gaude, E., Van Haute, L., Rebelo-Guiomar, P., Jackson, C.B., Rorbach, J., Pekalski, M.L., Robinson, A.J., Charpentier, M., Concordet, J.P., et al. (2016) Near-complete elimination of mutant mtDNA by iterative or dynamic dose-controlled treatment with mtZFNs. Nucleic Acids Res, 44, 7804–7816.

2. Gammage, P.A., Van Haute, L. and Minczuk, M. (2016) Engineered mtZFNs for Manipulation of Human Mitochondrial DNA Heteroplasmy. Methods Mol Biol, 1351, 145–162.

3. Nash, P.A., Turner, K.M., Powell, C.A., Van Haute, L., Silva-Pinheiro, P., Bubeck, F., Wiedtke, E., Marques, E., Ryan, D.G., Grimm, D., et al. (2025) Clinically translatable mitochondrial gene therapy in muscle using tandem mtZFN architecture. EMBO Mol Med, 17, 1222–1237.

4. Mok, B.Y., de Moraes, M.H., Zeng, J., Bosch, D.E., Kotrys, A.V., Raguram, A., Hsu, F., Radey, M.C., Peterson, S.B., Mootha, V.K., et al. (2020) A bacterial cytidine deaminase toxin enables CRISPR-free mitochondrial base editing. Nature, 583, 631–637.

5. Cho, S.I., Lee, S., Mok, Y.G., Lim, K., Lee, J., Lee, J.M., Chung, E. and Kim, J.S. (2022) Targeted A-to-G base editing in human mitochondrial DNA with programmable deaminases. Cell, 185, 1764–1776 e1712.

6. Yi, Z., Zhang, X., Tang, W., Yu, Y., Wei, X., Zhang, X. and Wei, W. (2024) Strand-selective base editing of human mitochondrial DNA using mitoBEs. Nat Biotechnol, 42, 498–509.

7. Zhang, X., Zhang, X., Ren, J., Li, J., Wei, X., Yu, Y., Yi, Z. and Wei, W. (2025) Precise modelling of mitochondrial diseases using optimized mitoBEs. Nature, 639, 735–745.

8. Mok, B.Y., Kotrys, A.V., Raguram, A., Huang, T.P., Mootha, V.K. and Liu, D.R. (2022) CRISPR-free base editors with enhanced activity and expanded targeting scope in mitochondrial and nuclear DNA. Nat Biotechnol, 40, 1378-1387.

9. Lee, S., Lee, H., Baek, G. and Kim, J.S. (2023) Precision mitochondrial DNA editing with high-fidelity DddA-derived base editors. Nat Biotechnol, 41, 378-386.

10. Xie, L., Cao, Y., Li, D., Ma, M., Jiao, D., Feng, H., Zuo, Z. and Zuo, E. (2024) Reducing off-target effects of DdCBEs by reversing amino acid charge near DNA interaction sites. Cell Res, 34, 877–881.

11. Lee, S., Lee, H., Baek, G., Namgung, E., Park, J.M., Kim, S., Hong, S. and Kim, J.S. (2022) Enhanced mitochondrial DNA editing in mice using nuclear-exported TALE-linked deaminases and nucleases. Genome Biol, 23, 211.

12. Silva-Pinheiro, P., Mutti, C.D., Van Haute, L., Powell, C.A., Nash, P.A., Turner, K. and Minczuk, M. (2023) A library of base editors for the precise ablation of all protein-coding genes in the mouse mitochondrial genome. Nat Biomed Eng, 7, 692-703.

13. Sabharwal, A., Kar, B., Restrepo-Castillo, S., Holmberg, S.R., Mathew, N.D., Kendall, B.L., Cotter, R.P., WareJoncas, Z., Seiler, C., Nakamaru-Ogiso, E. et al. (2021) The FusX TALE Base Editor (FusXTBE) for Rapid Mitochondrial DNA Programming of Human Cells In Vitro and Zebrafish Disease Models In Vivo. CRISPR J, 4, 799-821.

14. Guo, J., Zhang, X., Chen, X., Sun, H., Dai, Y., Wang, J., Qian, X., Tan, L., Lou, X. and Shen, B. (2021) Precision modeling of mitochondrial diseases in zebrafish via DdCBE-mediated mtDNA base editing. Cell Discov, 7, 78.

15. Qi, X., Chen, X., Guo, J., Zhang, X., Sun, H., Wang, J., Qian, X., Li, B., Tan, L., Yu, L. et al. (2021) Precision modeling of mitochondrial disease in rats via DdCBE-mediated mtDNA editing. Cell Discov, 7, 95.

16. Barrera-Paez, J.D., Bacman, S.R., Balla, T., Van Booven, D., Gannamedi, D.P., Stewart, J.B., Mok, B., Liu, D.R., Lombard, D.B., Griswold, A.J., et al. (2025) Correcting a pathogenic mitochondrial DNA mutation by base editing in mice. Sci Transl Med, 17, eadr0792.

17. Guo, J., Chen, X., Liu, Z., Sun, H., Zhou, Y., Dai, Y., Ma, Y., He, L., Qian, X., Wang, J. et al. (2022) DdCBE mediates efficient and inheritable modifications in mouse mitochondrial genome. Mol Ther Nucleic Acids, 27, 73–80.

18. Lee, H., Lee, S., Baek, G., Kim, A., Kang, B.C., Seo, H. and Kim, J.S. (2021) Mitochondrial DNA editing in mice with DddA-TALE fusion deaminases. Nat Commun, 12, 1190.

19. Silva-Pinheiro, P., Nash, P.A., Van Haute, L., Mutti, C.D., Turner, K. and Minczuk, M. (2022) In vivo mitochondrial base editing via adeno-associated viral delivery to mouse post-mitotic tissue. Nat Commun, 13, 750.

20. Kim, S., Kim, J., Cha, S., Ju, S., Lim, C.J., Hong, S., Bae, J., Oh, Y., Jung, S., Kim, S.P. et al. (2025) In vivo mitochondrial base editing restores genotype and visual function in a mouse model of LHON. Nat Commun, 16, 10923.

21. Metodiev, M.D., Spahr, H., Loguercio Polosa, P., Meharg, C., Becker, C., Altmueller, J., Habermann, B., Larsson, N.G. and Ruzzenente, B. (2014) NSUN4 is a dual function mitochondrial protein required for both methylation of 12S rRNA and coordination of mitoribosomal assembly. PLoS Genet, 10, e1004110.

22. Mutti, C.D., Van Haute, L. and Minczuk, M. (2024) The catalytic activity of methyltransferase METTL15 is dispensable for its role in mitochondrial ribosome biogenesis. RNA Biol, 21, 23–30.

