## Supplementary File 1 for "Development of the Mitochondrial Base Editor Analysis Package (MitoBEAP)"

### MitoBEAP

Lindsey Van Haute

2026-04-09

#### Introduction

MitoBEAP is an R package designed to analyze mitochondrial base editing data from gene editing experiments, both mitochondrial cytosine base editors (mitoCBE) such as DdCBE and adenine base editors (mitoABE) such as TALED. It calculates both on-target and off-target editing efficiencies, visualizes editing patterns, and produces reports for submission to data repositories.

The package is optimized for input from VarScan2 output in .csv format, and includes functionality for depth filtering, control heteroplasmy adjustment, heatmaps, and summary statistics.

For more detail, see our associated publication: "Development of the Mitochondrial Base Editor Analysis Package (MitoBEAP)." http://link.to.paper.com.

#### Input samples

This package is optimized to be used in combination with the output of VarScan2 (samtools mpileup -ABQ0 -aa –no-output-ins –no-output-ins –no-output-del –no-output-del –max-depth 0 –fasta-ref /path/to/ref.fasta | java -jar VarScan.jar readcounts –min-coverage 0) in .csv format. Example files can be downloaded here: https://github.com/NextGenSeek/MitoBEAP/ExampleFiles. However, other input files can be used too. We'll discuss this option later.

Before starting, ensure you are working in the correct directory: (setwd("/path/to/working/directory") ).

Then load the necessary libraries:

```
library(readr)
library(tools)
library(splitstackshape)
library(tibble)
library(ggplot2)
library(dplyr)
library(tidyr)
library(ggrepel)
library(RColorBrewer)
library(MitoBEAP); packageVersion("MitoBEAP")
```

```
## [1] '0.1.0'
```

For this tutorial, we're using files included with the package. If you want to use your own files, you can upload your own files.

If the coverage of a region is too low, a potential single read error will result in high "off-target percentages". Therefore, MitoBEAP requires the user to define the minimum depth or coverage you want to include. We would recommend not to include positions below a depth of 100. Depending on your experiment or amplicon choice, you might also be interested in a certain region only.

To begin, list your raw VarScan2 output files and define a minimum depth threshold for filtering. Skip this step if you have a different type of input files:

```
# List input files
RawCountFiles <- list.files("./raw",
                            pattern = "*.csv",
                            full.names = TRUE)

# Specify prefered minimum depth first
depth = 100

# Run the following command
lapply(RawCountFiles, LoadRawCount)

# To select a certain region use: this can be done with the function RegionSelection:

RegionSelection(RawCountFiles) # Default range: from start to end
RegionSelection(RawCountFiles, position_range = c(1000, 5000), depth_filter = TRUE, min_depth = 100)
```

To use all functions of the package, the user needs to generate a SampleList or upload a .csv with sample information FileName (file name, without .csv), SampleName (how you want the samples to be labelled) and Condition (e.g. control - treated) as shown in the example and name this SampleList.

```
SampleList <- read.delim("filename.csv",
                         row.names = NULL,
                         sep = ",")
head(SampleList)
```

```
##   Order SampleName FileName Condition Replicate
## 1     3      12S-1   E8_S56   treated         1
## 2     4      12S-2   E9_S57   treated         1
## 3     5      12S-3  E10_S58   treated         2
## 4     6      12S-4  E11_S59   treated         3
## 5     7      12S-5  E12_S60   treated         1
## 6     8      12S-6   F1_S61   treated         1
```

#### Run Base Editing Analysis

Next, we'll perform the basic calculations. For this step, there are two variants, depending on the input files (VarScan2 as in the example) or alternative input files. As this tool is specifically designed to analyse cytosine base editors (CBE) or adenine base editors (ABE), it will only take C-T and G-A (CBE) or A-T to G-C (ABE) conversions into account and will ignore other changes, as those are existing variants or technical errors. In this step, you also have to assign the on-target position for your analysis. This matches the position of the reference genome you used to create the original input files.

The function below can be used if you use VarScan2 input files, like in our example. Otherwise, use the subsequent code (FilterCGEdits/FilterATEdits).

```r
# List files. If you didn't change anything, you can just use the line below:
MinXFiles <- list.files("./minimum",full.names = TRUE) #Is there a way to avoid this line?

# On-target position:
OntargetPosition <- 1561 #CHANGE ME

# Run script for CBE
lapply(MinXFiles, OnTargetCalcCBE)
# Alternatively, run script for ABE
lapply(MinXFiles, OnTargetCalcABE)
```

The following shunk should NOT be run if you follow the VarScan2 script. If you have different input files, make sure they're in the correct format. MitoBEAP expects .csv files with the following columns: chrom, position, ref_base, depth, base (= alternative base), reads, percentage.

```r
# Select input files
allFiles <- list.files("./folder", pattern = "\\.csv$", full.names = TRUE)

# Run calculator:
FilterCGEdits(allFiles) #CBE
# or
FilterATEdits(allFiles) #ABE
```

From here onwards, all steps should be the same, regardless of the input files.

#### Summary Table Generation

Use the following functions to generate overall summaries of coverage, on-target, and off-target activity:

```r
# If you didn't change anything, you can use the lines below
CoverageFiles <- list.files("./Coverage",
                            pattern = "*.txt",
                            full.names = TRUE)
Coverage <- as.data.frame(CoverageCalc(CoverageFiles))

MeanFiles <- list.files("./mean", pattern = "*mean.txt",
                        full.names = TRUE)
Mean <- as.data.frame(MeanCalc(MeanFiles))

OnTargetFiles <- list.files("./OnTarget", pattern = "*ontarget.txt",
                             full.names = TRUE)
OnTarget <- as.data.frame(CombineOnTarget(OnTargetFiles))

CreateOverview()
```

This creates All_Ontarget_mean_Coverage.csv in the ./Overview directory.

```
##    Sample        Off.target.. On.target.. Coverage RealName Order
## 1 E10_S58 1 0.557623497639187    8.970727 1903.158    12S-3     6
## 2 E11_S59 1 0.580606419927107   14.425646 2094.053    12S-4     5
## 3 E12_S60 1 0.551971175658703   41.020236 2344.339    12S-5     4
## 4  E8_S56 1 0.691491938866253    9.717097 1485.268    12S-1     8
```

```
## 5  E9_S57 1 0.634058456164951   48.805879 1540.712    12S-2     7
## 6  F1_S61 1 0.984706934847988   39.694656  891.073    12S-6     3
## 7  F2_S62  1 0.71870789707064   16.525424 1336.088    12S-7     2
## 8  F3_S63 1 0.504355763258661   40.133779 2463.899    12S-8     1
```

#### Export Excel Workbook

MitoBEAP also has the option to generate an excel spreadsheet with all data, which could be used as overview table when uploading the raw fastq files to a data repository. A spreadsheet can be easily created by the following command:

```
CreateSpreadsheet()
```

After running this command, the spreadsheet should now be available in the folder Overview

#### Depth and Heteroplasmy Overview

Some users might also find the following file useful: The function DepthFile generates a .csv file that shows the heteroplasmy percentage as well as the coverage depth for each position and each sample. Default settings show the entire genome, but users can specify the region of interest. The generated file can also be used to generate heatmaps or other figures using excel, Prism or other software packages.

```
DepthFile() # default is from start to end

# DepthFile(fromP = 1, toP = 500) # generates the file for the first 500 positions of the genome
```

#### Visualization options

MitoDD also has several options to easily visualize the samples. One of the simples figures is a barchart to show the on-target base editing for each sample:

```
CreateBarChart(data_type = "OnTarget", colour = "lightgreen")
```

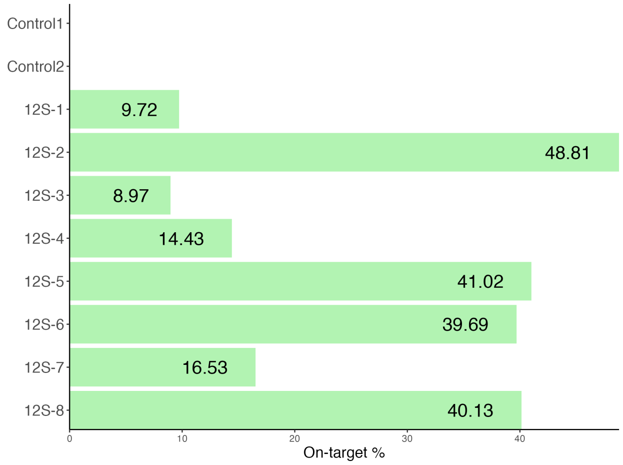


Or a barchart showing the mean off-target levels

```
CreateBarChart(data_type = "OffTarget", colour = "skyblue")
```

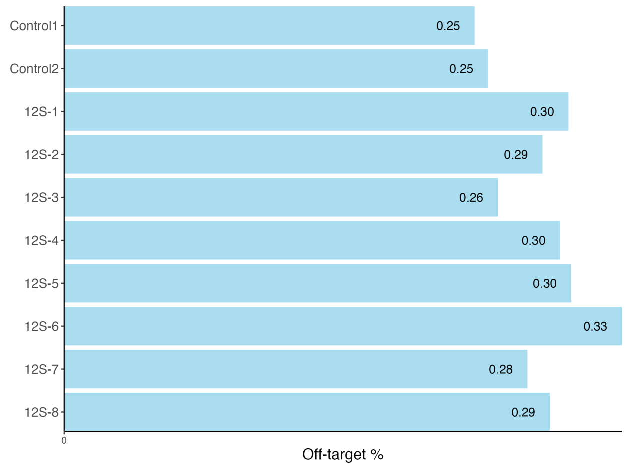


#### Scatterplots of Off-target Editing

To visualize off-target effects for each sample individually, scatter plots are very helpful. Again, these can easily be generated. The user can specify to label the dots above a certain level by position.

```
DdCBE_df()
AdjScatter(labelPercentage = 10) # Default = 10
```

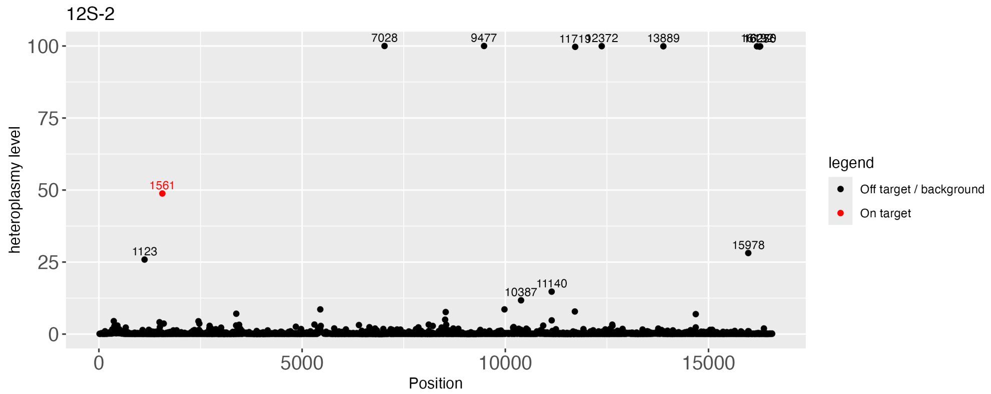


#### Remove Control Backgrounds (heteroplasmy)

For most experiments, the cells or tissue analysed, will not 100% match with the reference genome. It is expected that these positions remain unaltered during the experiment, so they could be considered background. However, the user might want to remove these positions, both for visualization and off-target calculation purposes. Therefore, the user can specify to remove positions that have more than X % heteroplasmy levels in some or all of the control samples. If the user prefers not to remove those highly heteroplasmic positions, max_threshold = 100 can be used

The user can also further optimize the figure, by removing dots below a certain value from the figure (however, it should be noted that those positions will be taken into account for off-target calculations)

To exclude polymorphic positions present in controls:

```
DdCBE_df(max_threshold = 90, controls = 3)
AdjScatter(min_threshold = 1.2, labelPercentage = 10) # labelPercentage = default = 10
```

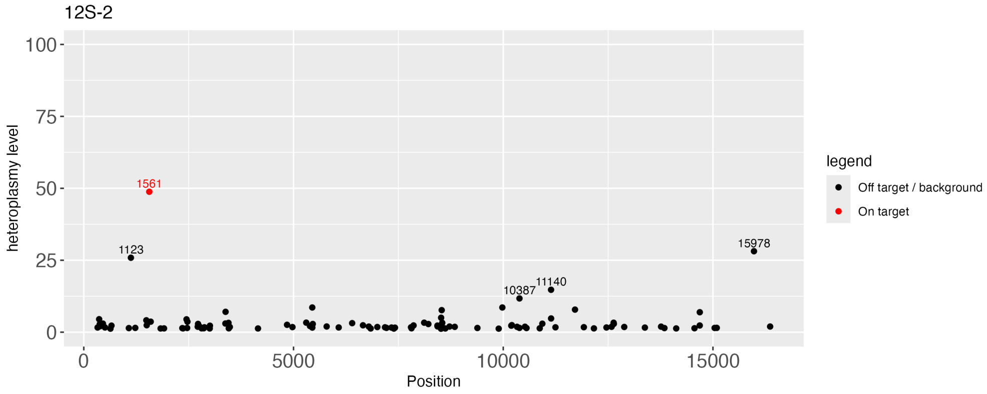


The user might also want to recalculate off-target averages and create a barchart with the above specified maximum heteroplasmy levels in controls.

```
# Recalculate off-target average with this max_threshold
AdjOffTarget()
# Create barchart with this max_threshold
AdjCreateBarChart(colour = "skyblue")
```

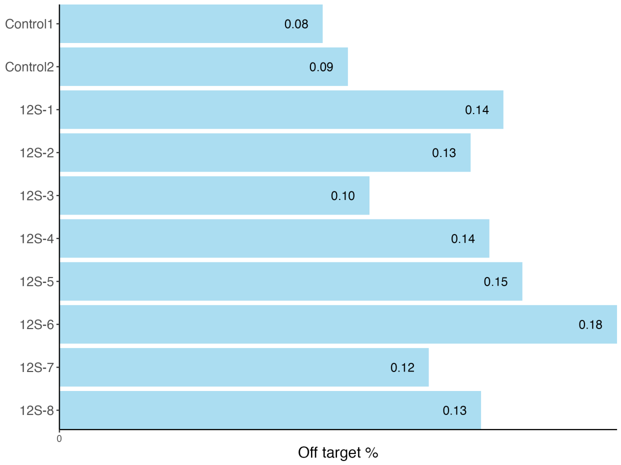


#### Comparative Plot: On- vs Off-target

This plot shows on-target versus adjusted off-target editing:

```
OnVsOffTargetAdj() #standard = Condition = TRUE, FALSE= no condition information
# default xlab = "Off target effects (%)",
# default ylab = "Heteroplasmy level (%)"
# default ggtitle = "On-versus off-target effects"
```

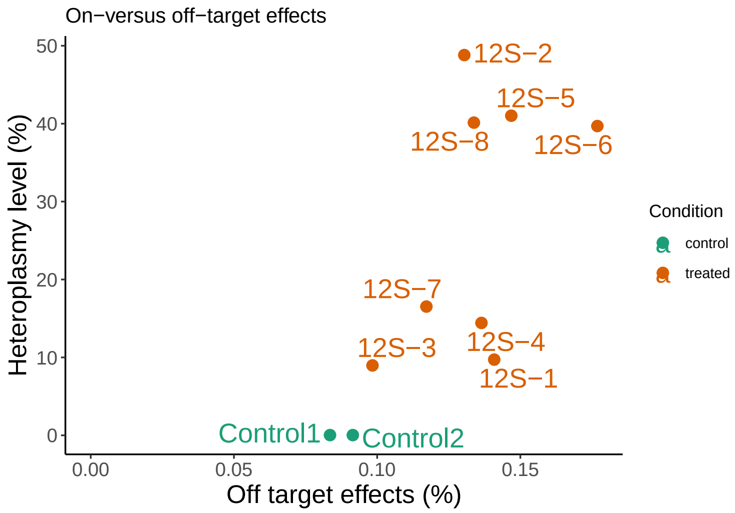


#### Comparative Plot: grouped results

This function generates a bar chart showing the mean of the replicates with error bars (sd).

In order to use this function, the SampleList should contain a column "CombinedName" with the group names.

```
off_target_count_summary()
# default lower = 0.1
# default upper = 70
# default condition_levels = c("control", "treated")
# default out_dir = "."
# default csv_prefix = "MutationCount"
# default plot_file = "MutationCount_grouped.pdf"
```

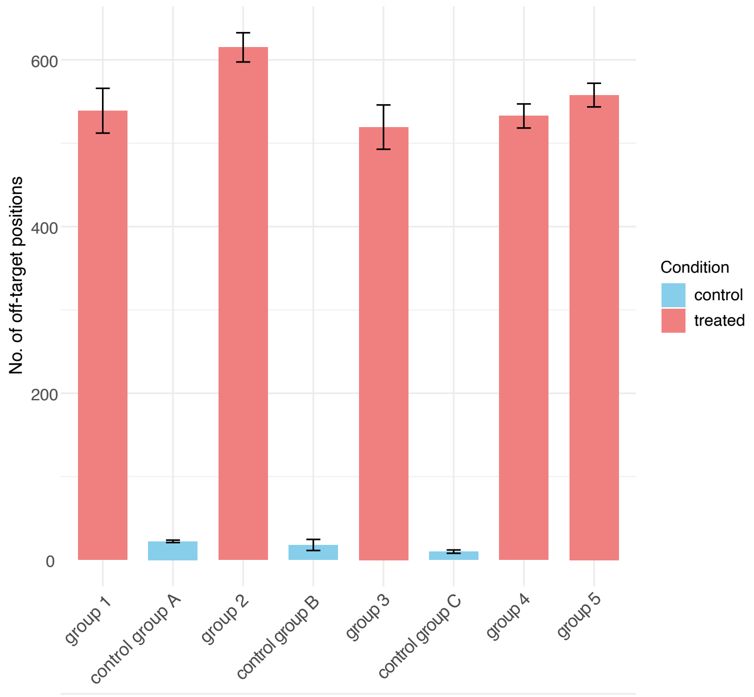


#### Heatmap of Bystander Editing

Base editing can be very specific, but bystander mutations can often be detected in the activity window of the base editor, so this requires extra attention.

Visualize editing levels near the on-target position:

```
# BystanderDistance()

Bystander(BystanderDistance = 10,
          title = "Bystander effect",
          x= "mtDNA position",
          y= NONE)
```

This creates the desired heatmap in the folder Plots

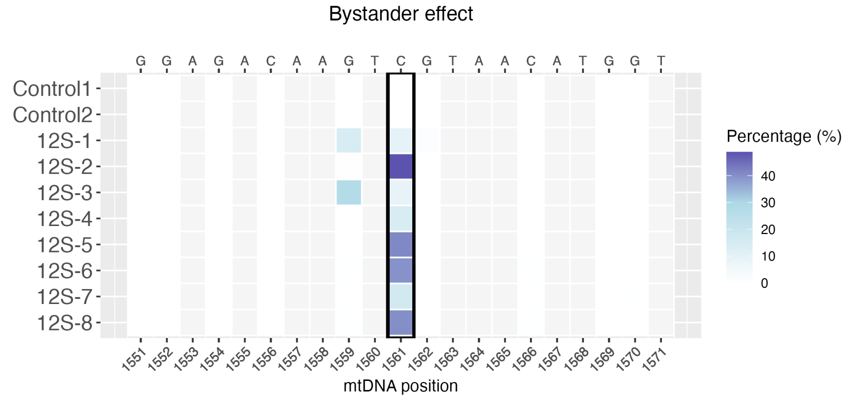


Alternatively, to create this heatmap with only the potential target sites (CG for CBE or AT for ABE) use the following function

```
# BystanderDistance()

bystander_only_relevant(BystanderDistance = 10,
                        title = "Bystander effect",
                        x= "mtDNA position",
                        y= NONE)
```

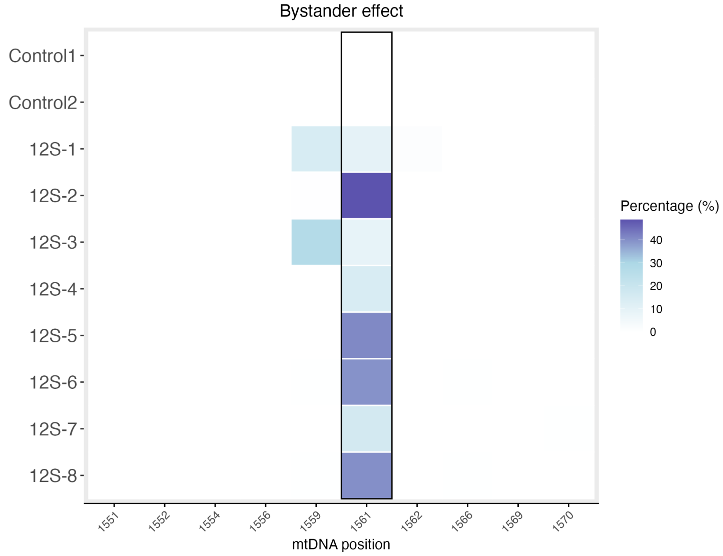

#### Histogram of off-target effect

```
plot_OffTarget_histograms(bw = 0.1, min_pct = 0.1, max_pct = 5)
# default binwidth bw = 1
# default min_pct = 0
# default max_pct = Inf
```

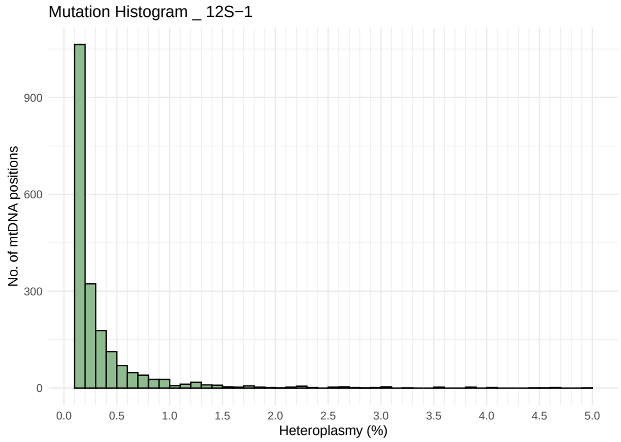


#### Conclusion

MitoBEAP offers a fast and reproducible way to analyze mitochondrial base editing datasets, with robust filtering and visualization options. It is adaptable to various types of experiments and ready for integration with data repositories.

For questions or contributions, please visit our GitHub page. https://github.com/NextGenSeek/MitoBEAP
