## Supplementary figures and images for "Development of the Mitochondrial Base Editor Analysis Package (MitoBEAP)"

### Supplementary Figure 1

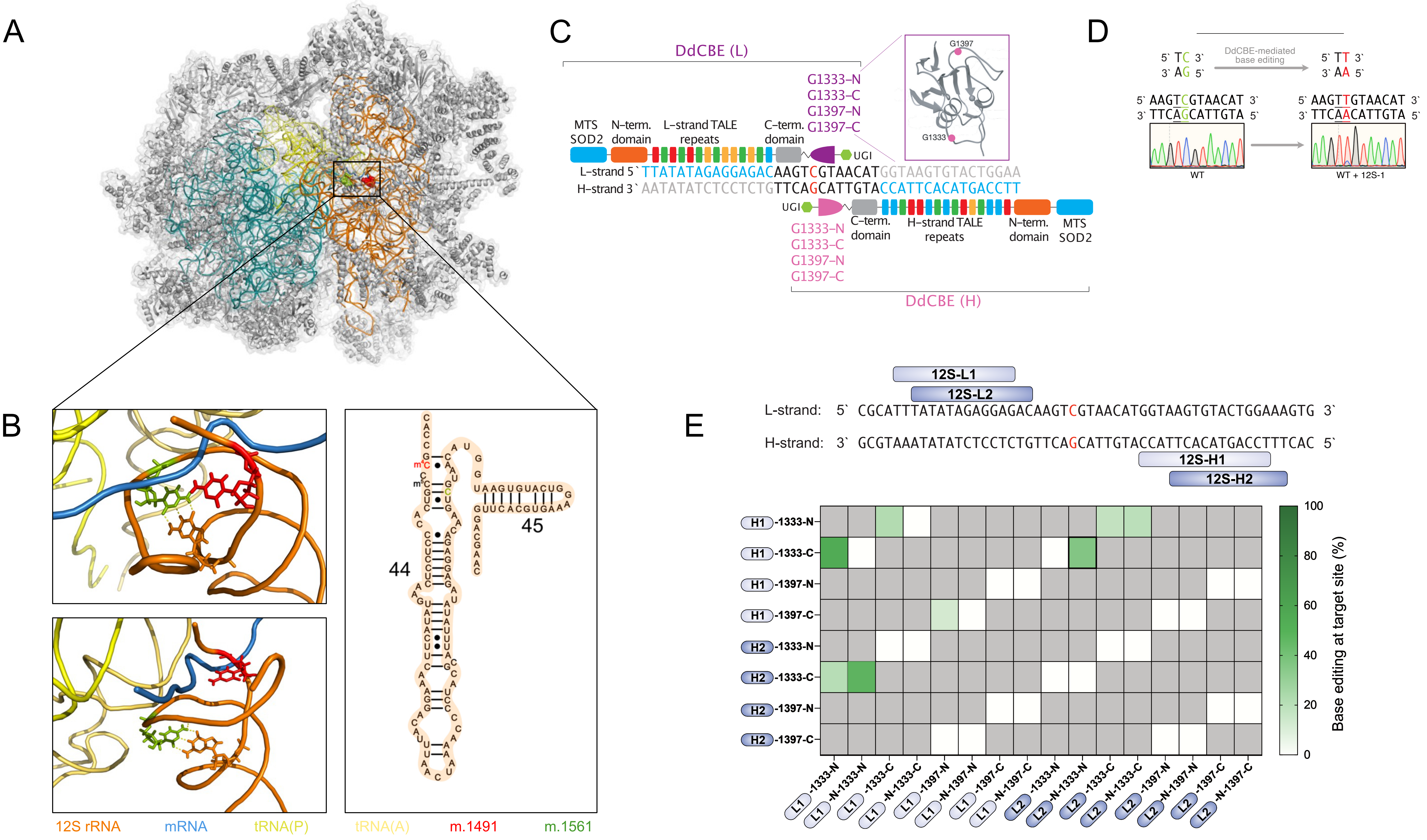

### Supplementary Figure 2

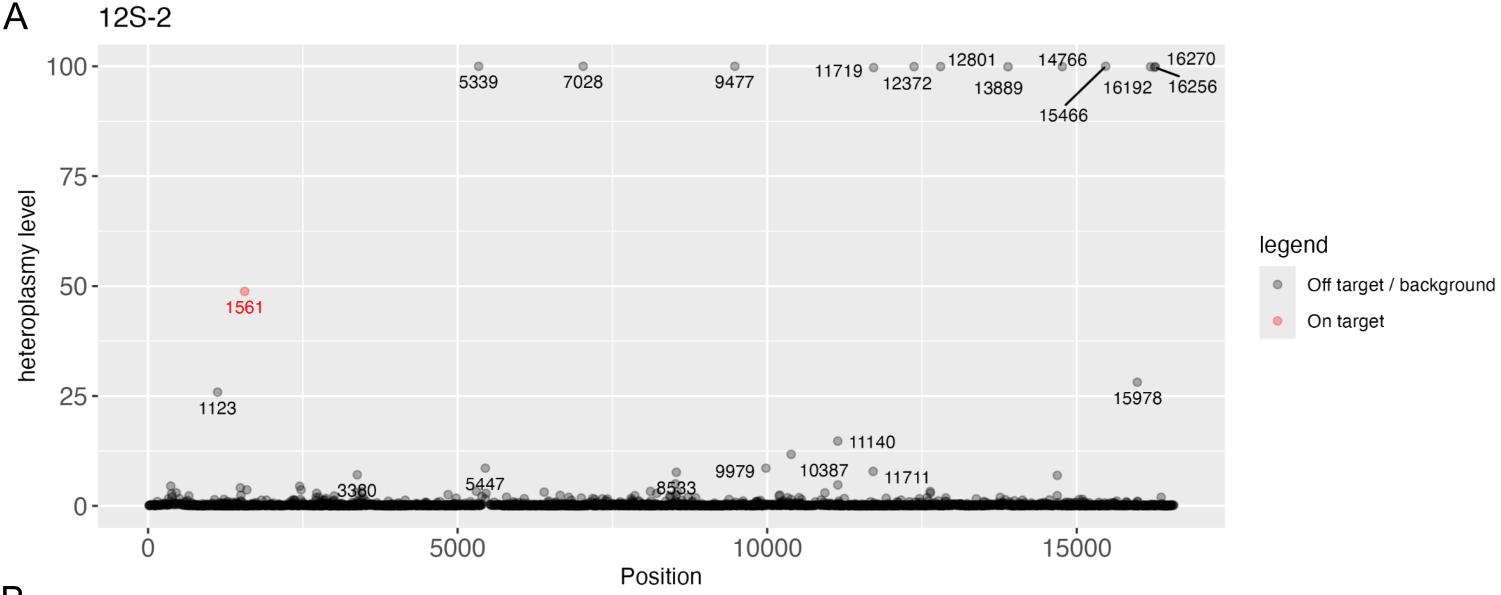

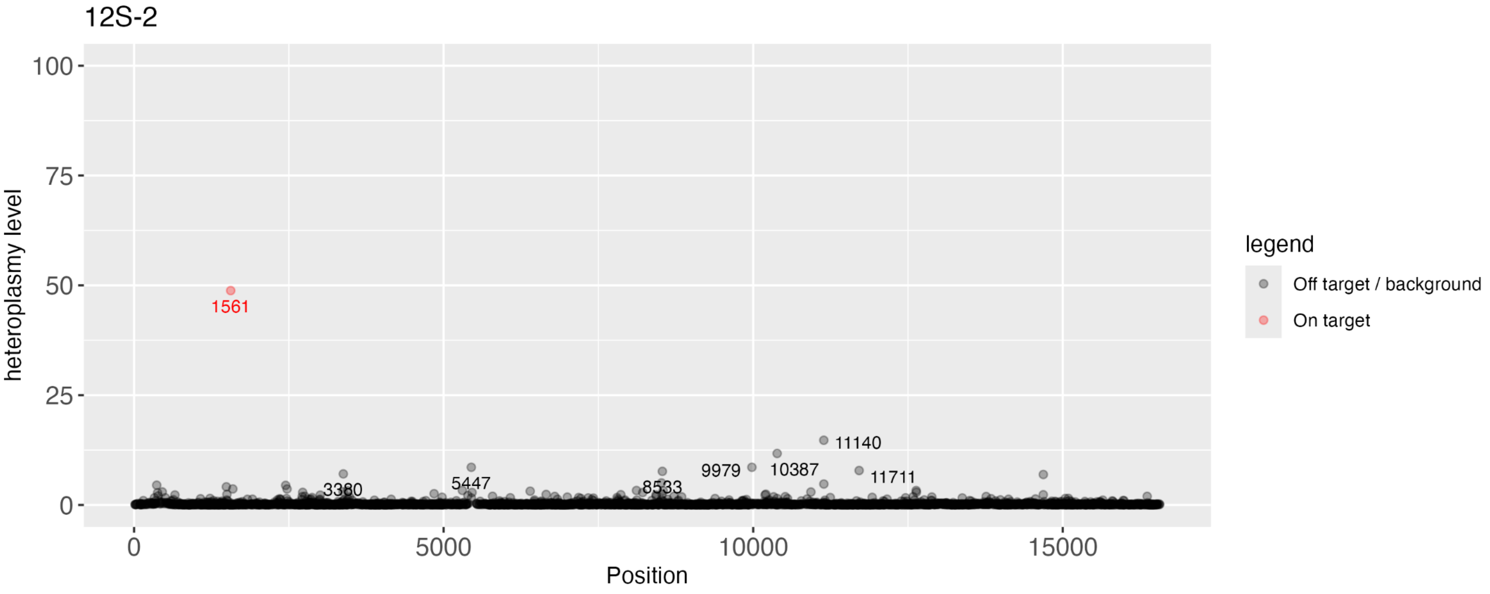

### Supplementary Figure 3

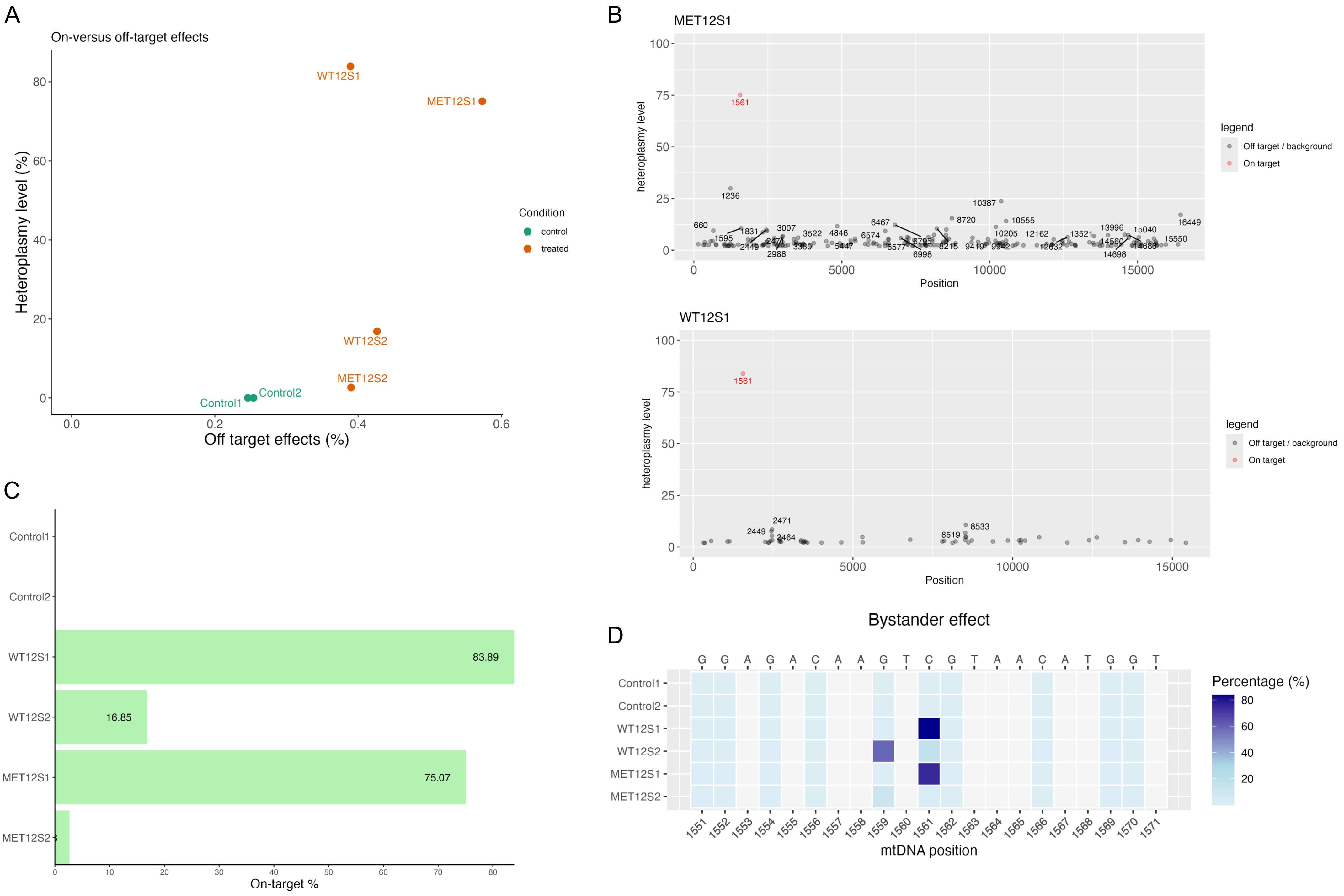
